# A deep learning-based toolkit for 3D nuclei segmentation and quantitative analysis in cellular and tissue context

**DOI:** 10.1101/2024.02.19.580954

**Authors:** Athul Vijayan, Tejasvinee Atul Mody, Qin Yu, Adrian Wolny, Lorenzo Cerrone, Soeren Strauss, Miltos Tsiantis, Richard S. Smith, Fred A. Hamprecht, Anna Kreshuk, Kay Schneitz

## Abstract

We present a new set of computational tools that enable accurate and widely applicable 3D segmentation of nuclei in various 3D digital organs. We developed a novel approach for ground truth generation and iterative training of 3D nuclear segmentation models, which we applied to popular CellPose, PlantSeg, and StarDist algorithms. We provide two high-quality models trained on plant nuclei that enable 3D segmentation of nuclei in datasets obtained from fixed or live samples, acquired from different plant and animal tissues, and stained with various nuclear stains or fluorescent protein-based nuclear reporters. We also share a diverse high-quality training dataset of about 10,000 nuclei. Furthermore, we advanced the MorphoGraphX analysis and visualization software by, among other things, providing a method for linking 3D segmented nuclei to their surrounding cells in 3D digital organs. We found that the nuclear-to-cell volume ratio varies between different ovule tissues and during the development of a tissue. Finally, we extended the PlantSeg 3D segmentation pipeline with a proofreading script that uses 3D segmented nuclei as seeds to correct cell segmentation errors in difficult-to-segment tissues.

**Summary Statement:** We present computational tools that allow versatile and accurate 3D nuclear segmentation in plant organs, enable the analysis of cell-nucleus geometric relationships, and improve the accuracy of 3D cell segmentation.

## Introduction

Tissue morphogenesis is a complex, multi-scale process that ultimately results in an organ or tissue of a specific size and shape and characteristic 3D cellular architecture. Advances in imaging increasingly allow generation of 3D digital organs with cellular resolution, which are useful tools for unraveling the integration and feedback processes between molecular regulatory circuits and the cellular architecture of developing tissues and organs. Plants are excellent systems for generating 3D digital organs because their cells are immobile and the cellular architecture of plant organs can be easily observed using various types of microscopy.

Over the years, and partly through the application of artificial intelligence, powerful open-source software packages have been developed for 3D cell segmentation of confocal microscopy images (Barbier de Reuille et al., 2015; Eschweiler et al., 2019; Fernandez et al., 2010; Schmidt et al., 2014; Sommer et al., 2011; Stegmaier et al., 2016). Machine learning based software, including CellPose, PlantSeg and StarDist, represents a recent advance in this area, providing improved 3D segmentation of tissues at cellular resolution (Eschweiler et al., 2019; Stringer et al., 2021; Weigert et al., 2020; Wolny et al., 2020). The output of such pipelines can then be quantitatively analyzed in image analysis software like MorphoGraphX (Barbier de Reuille et al., 2015; Strauss et al., 2022). The advances in these computational resources have enabled the generation of a number of digital 3D models of a variety of plant organs, which have allowed single-cell analysis in 3D and have been instrumental in gaining fundamental insights into various processes in plants, including embryo, root, and ovule development (Bassel et al., 2014; Fridman et al., 2021; Graeff et al., 2021; Hernandez-Lagana et al., 2021; Lora et al., 2017; Montenegro-Johnson et al., 2015; Ouedraogo et al., 2023; Pasternak et al., 2017; Schmidt et al., 2014; Vijayan et al., 2021; Yoshida et al., 2014).

An important feature that is presently missing from these 3D digital models is the integration of the size and shape of the nuclei into the cellular framework. The ability to not only robustly segment nuclei in 3D, even in deeper tissues, but also to link the 3D architectures of nuclei and their surrounding cells in a tissue-specific context enables the study of central biological processes such as nuclear size control (Cantwell and Nurse, 2019c). Another key process is the control of gene expression. Spatial gene expression patterns as well as expression levels can be assessed with cellular resolution, for example, using ratiometric nuclear reporters driven by gene-specific promoters (Federici et al., 2012).

ClearSee-based protocols for cleared whole-mount preparations of plant organs allow staining of cell walls and nuclei with various cytological dyes without the need for transgenic plants carrying the appropriate reporter constructs and maintain compatibility with reporters based on fluorescent proteins (Kurihara et al., 2015; Musielak et al., 2015; Tofanelli et al., 2019; Ursache et al., 2018). The establishment of the 3D digital reference atlas of Arabidopsis ovule development represents a recent example that used this approach (Vijayan et al., 2021). During the preparation of the atlas, ovules were fixed and cleared with ClearSee (Kurihara et al., 2015). Cell outlines were stained with the cell wall stain SCRI Renaissance (SR2200) (Harris et al., 2002; Musielak et al., 2015), while the nuclei were stained with TO-PRO-3 (Bink et al., 2001; Van Hooijdonk et al., 1994). The digital ovule atlas provided detailed insight into the 3D cellular architecture of the ovule but lacked information on the size and shape of the nuclei. TO-PRO-3 stains double-stranded nucleic acids and can therefore be a useful tool for 3D volumetric nuclear extraction. However, the signal intensity of any typical nuclear stain can exhibit variable intensities, scatter, and photobleaching when imaging deeper tissue layers, rendering accurate 3D nuclear segmentation extremely difficult.

Therefore, our overall goal is to accurately segment plant nuclei in 3D images with weakly stained nuclei. Several deep learning-based segmentation algorithms have recently been proposed for this task: PlantSeg (Wolny et al., 2020), Cellpose (Stringer et al., 2021), and StarDist (Weigert et al., 2020). However, none of them can be used out of the box. PlantSeg and CellPose pre-trained models have not been exposed to weakly stained plant nuclei while 3D StarDist does not provide trained models and requires retraining. The main bottleneck for model training is the lack of publicly available 3D ground truth with correctly delineated nuclei. This step is famously labor-intensive even for high-contrast, high signal-to-noise ratio (SNR) image volumes.

In this study, we combine different staining strategies to quickly achieve 3D segmentation ground truth for model training. Together with human-in-the-loop correction, we use this approach to acquire fully annotated volumes of weakly stained nuclei. On this basis, we train highly accurate segmentation networks, which we show to be generalizable to other datasets obtained by various imaging methods and from a variety of plant and animal tissues labeled with different staining methods. In addition, we introduce a combination of processes in MorphoGraphX that associates each nucleus with the cell in which it resides, and that provides the nucleus with the cells’ respective tissue labels. It allows the investigation of various cell-nucleus relationships, such as the nucleus-to-cell volume (N/C) ratio. We demonstrate the general value and broad applicability of these technical advances in proof-of-concept analyses.

## Results

### A novel iterative approach to ground truth generation and 3D nuclear model training

In a first attempt at 3D nuclear segmentation of TO-PRO-3-stained ovule nuclei in 3D image stacks, we found that the available plant nuclei segmentation model in PlantSeg did not yield segmented nuclei of sufficient quality for ground truth generation. Thus, we employed Cellpose (Pachitariu and Stringer, 2022; Stringer et al., 2021) as it had an existing nuclei model used for 3D nuclear segmentation. However, we still observed improper segmentation with errors in detecting and separating nuclear borders (Fig. 1A-D). This is probably due to the TO-PRO-3 nuclear staining being variable and often quite weak and diffuse, particularly in deeper layers. In addition, the signal was absent in the nucleolus, resulting in an uneven nuclear surface and segmentation that looked like a hole extruded from the nuclear surface (Fig. 1C).

**Fig. 1.**
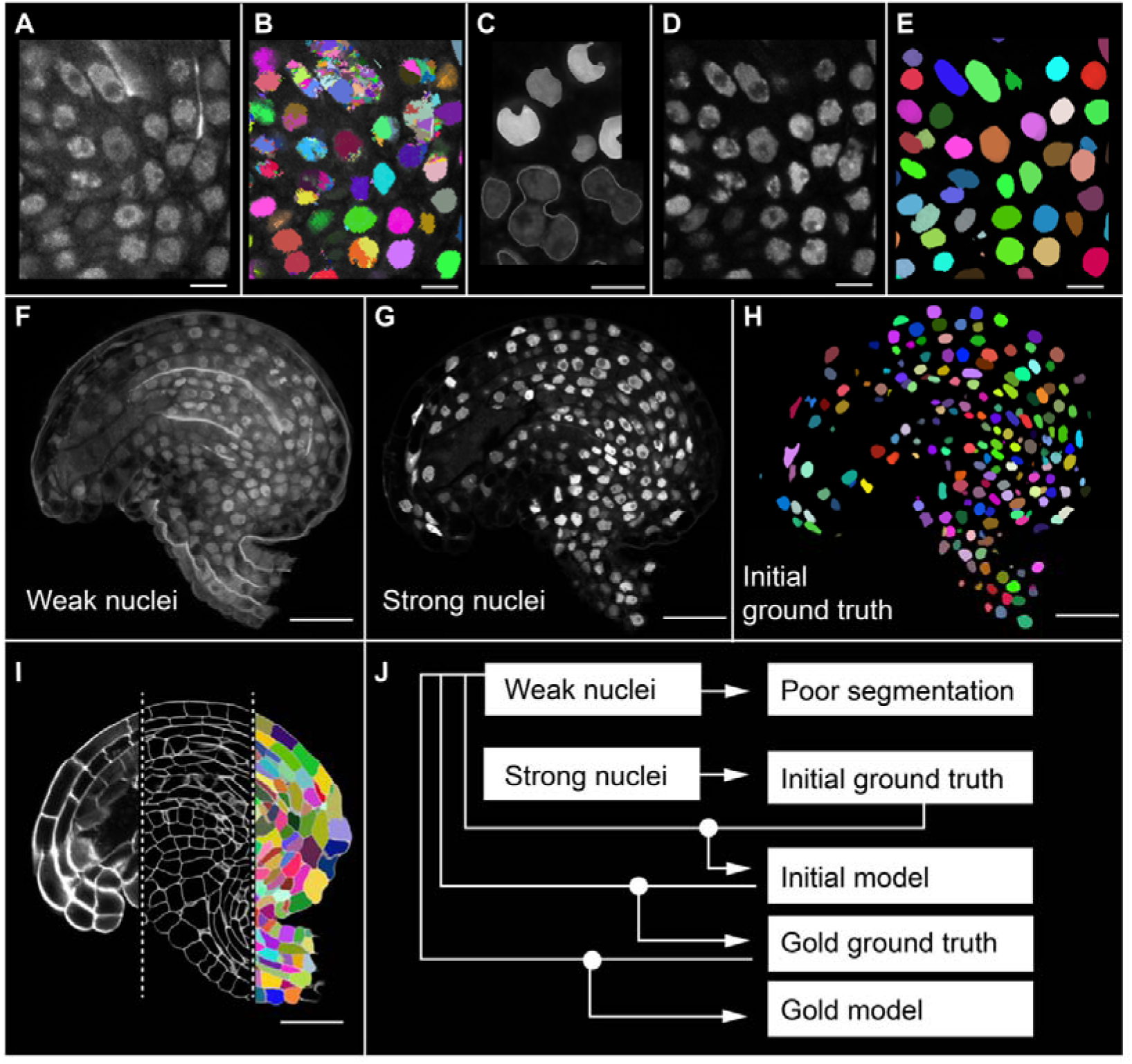
3D dataset for model training. (A) 2D section view of TO-PRO-3-stained nuclei in Arabidopsis ovules. (B) 3D nuclear segmentation of weak nuclei stain performed using Cellpose nuclei model. (C) A zoomed-in view displaying the erroneous segmentation. Typical segmentation errors in the nuclei stains segmentation resulting in improper size, shape and number of nuclei. (D) Fluorescent nuclei reporter H2B: tdTomato raw image. (E) 3D Cellpose nuclei model segmentation of the bright tdTomato nuclei fluorescence. (F-I) 2D section view from one of the five training dataset. (F) Weak nuclei channel (TO-PRO-3-stained) used for training. (G) Strong nuclei channel (nuclei reporter H2B: tdTomato) used for generating ground truths. (H) Initial ground truth used for training initial model. 3D nuclear segmentation of the strong nuclei channel performed using the Cellpose nuclei model. (I) Raw cell wall stain, PlantSeg cell boundary predictions and cell segmentation available with the training dataset (from left to right) (J) Illustration of model training strategy. Scale bars: 5µm (A-E); 20 µm (F-I).

To address these issues, we developed a novel strategy based on samples that simultaneously show strong and faint signals in the nuclei that can be collected in separate channels. We first generated a transgenic line expressing a translational fusion of the fluorescent protein tdTomato to histone H2B driven by the *UBIQUITIN10* (*UBQ*) promoter (pUBQ::H2B:tdTomato). Ovules of this reporter line were fixed, cleared, and stained with the cell wall stain SR2200 and the nuclear stain TO-PRO-3. Ovules were imaged and the SR2200, TO-PRO-3, and H2B:tdTomato signals were collected in three separate channels (Fig. 1F,G,I). The broadly expressing nuclear pUBQ::H2B:tdTomato reporter provided a strong and uniform nuclear signal that could be segmented into nuclei using the standard Cellpose nuclear model (Fig. 1D,E,G,H). We then used the results of human proofread instance nuclear segmentation of the strong H2B:tdTomato reporter channel as the “initial ground truth” for training three sets of initial 3D models: PlantSeg_3Dnuc_initial, StarDist-ResNet_3Dnuc_initial, and Cellpose-Finetune-nuclei_3Dnuc_initial. The PlantSeg and StarDist initial models were trained on the weak TO-PRO-3 nuclear stain channel using the neural networks implemented in the respective pipelines. The Cellpose initial models were trained on the TO-PRO-3 channel by fine-tuning the pretrained Cellpose “nuclei’’ model. The segmentation results using the initial models turned out to be still imperfect and required several corrections by an expert.

To obtain further model improvements we applied an iterative training strategy (Fig. 1J). We used the StarDist-ResNet_3Dnuc_initial model to segment the original weak TO-PRO-3-based nuclear stain channel as it provided the best qualitative results, resulting in a modified ground truth. This modified ground truth was then human proofread, resulting in the “gold ground truth”. In a next step, the “gold ground truth” and the original weak TO-PRO-3-based nuclei stain were used to train six sets of 3D “gold models’’ using one or multiple neural networks implemented in PlantSeg, Cellpose, and StarDist (Table 2), probing for the best parameter settings.

We tested how much model performance improved when human-in-the-loop (HITL) was involved, i.e., initial vs gold model. To this end we employed a quantitative comparison of initial and gold PlantSeg, StarDist-ResNet and Cellpose-Finetune-nuclei models. We made use of the imperfect initial models to generate modified and better ground truth by involving a HITL proofreading before using them for the training that resulted in the “gold models’’. (Fig. 1J). The detailed description of model training including the datasets used for training and testing are provided in the “Model training and score quantification” section of Materials and Methods. Comparison of model performance between initial and gold PlantSeg, Cellpose-Finetune-nuclei and StarDist-ResNet models was performed by 5-fold average precision (AP) score quantification (Table 1). Results indicate that all methods demonstrate increased performance after gold training. PlantSeg and StarDist-ResNet gold models turned out to be superior to the Cellpose-Finetune-nuclei gold models and demonstrated high precision segmentation compared to the respective initial models.

**Table 1.** Comparative analysis of different model performance when involving human in the loop to train a gold model.

| Tool used | Model | Version | 5-fold AP ± STD |
| --- | --- | --- | --- |
| PlantSeg | UNet GASP | Initial | 57.40% ± 7.7% |
| PlantSeg | UNet GASP | gold | 78.80% ± 1.98% |
| StarDist | ResNet | Initial | 67.61% ± 6.5% |
| StarDist | ResNet | gold | 78.33% ± 1.73% |
| Cellpose | Finetune nuclei | initial | 43.64% ± 12.88% |
| Cellpose | Finetune nuclei | gold | 51.96% ± 12.51% |

Segmentation of the test dataset was performed using each of the listed initial and gold models and the mean average precision is scored for different methods compared to gold ground truth.

### Comparisons of the different gold models

Quantitative and qualitative performance comparisons of the different gold models were performed and results are presented in Table 2, Fig. 2, and Fig. S1. With the exception of the Cellpose-derived models, all other gold models performed excellently on the raw images of nuclei stains as can be seen with qualitative comparison (Fig. 2, Fig. S1). The weak nuclei signals were strongly detected especially with the proposed new PlantSeg_3Dnuc_gold, StarDist-ResNet_3Dnuc_gold and Stardist-UNet_3Dnuc_gold models. Segmented nuclei surfaces were devoid of any artifacts like an extruded hole as in the raw nuclei image segmentation prior to developing this method. The AP scores obtained in these cases were very high when compared to the proposed new Cellpose nuclei gold models (Table 2). Average precision graphs also clearly indicate high precision of the PlantSeg, StarDist-ResNet, and StarDist-UNet gold models and how little they vary compared to the Cellpose gold models (Fig. S1I-N).

**Table 2.** Comparative analysis of different gold model training performance.

| Tool used | Model | 5-fold AP ± STD |
| --- | --- | --- |
| PlantSeg | UNet GASP | 78.80% ± 1.98% |
| StarDist | ResNet | 78.33% ± 1.73% |
| StarDist | UNet | 78.25% ± 1.84% |
| Cellpose | Finetune nuclei | 51.96% ± 12.51% |
| Cellpose | Finetune cyto2 | 51.05% ± 12.93% |
| Cellpose | Trained from scratch | 51.26% ± 13.75% |

Segmentation of the test dataset was performed using each of the listed methods and the mean average precision is scored for different methods compared to a human proofread ground truth. The configuration files used for training can be found along with the model.

**Fig. 2.**
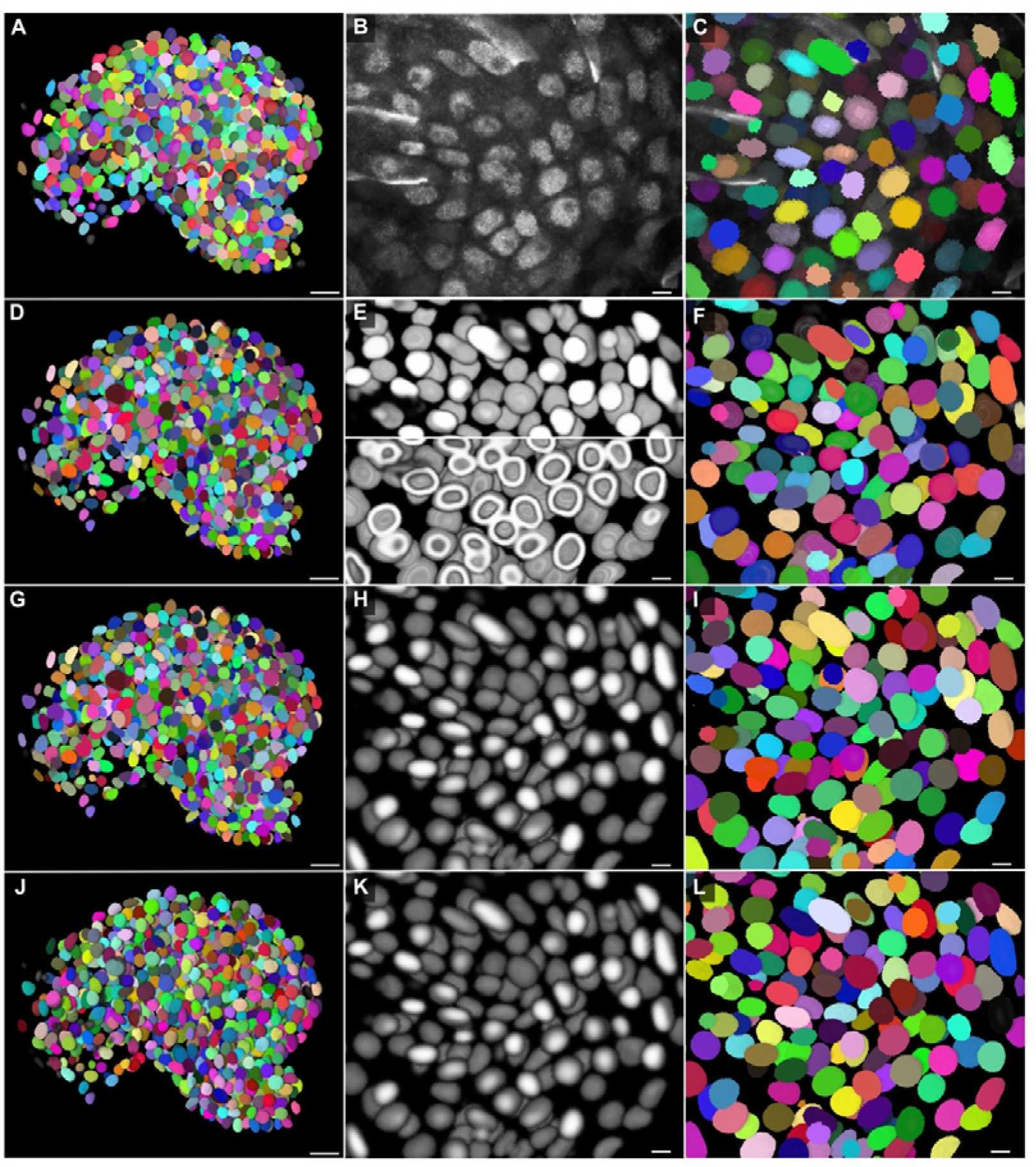
Qualitative comparison of segmentation results using different trained models. Qualitative comparison displaying the Arabidopsis ovule testing dataset 1135 (N5 dataset) with trained model (Model-5) using four other training datasets. (A) 3D view of ground truth nuclear segmentation. (B) Zoomed 2D section view of raw weak TO-PRO-3 iodide nuclei stain. (C) Ground truth nuclear segmentation corresponding to the zoomed view in (B). (D-E) PlantSeg predictions and segmentation using the proposed PlantSeg model. (D) 3D PlantSeg GASP segmentation performed using the proposed PlantSeg model. (E) View corresponding to (B) showing PlantSeg nuclei predictions. Top panel: PlantSeg nuclei center predictions. Bottom panel: PlantSeg nuclei envelope prediction from raw data. (F) PlantSeg GASP segmentation of the corresponding section in (B). (G-I) StarDist ResNet nuclei predictions and segmentation using the proposed ResNet model. (G) StarDist ResNet 3D nuclear segmentation performed using the proposed StarDist model. (H) View corresponding to (B) showing StarDist ResNet nuclei predictions. (I) StarDist ResNet nuclear segmentation of the corresponding section in (B). (J-L) StarDist UNet nuclei predictions and segmentation using the proposed UNet model. (J) StarDist UNet 3D nuclear segmentation performed using the proposed StarDist model. (K) View corresponding to (B) showing StarDist UNet nuclei predictions. (I) StarDist UNet nuclear segmentation of the corresponding section in (B). Scale bars: 10μm.

### StarDist-ResNet and PlantSeg gold models are two highly reliable models

The PlantSeg model was trained to produce a nuclear center probability map and a nuclear envelope probability map (Fig. 2E). The nuclear envelope probability was processed by Generalized Algorithm for Signed Graph Partitioning (GASP) (Bailoni et al., 2019) to obtain an initial instance segmentation, which is then filtered according to the probability of the nuclei center (Fig. 2D,F). Data volumes for both training and inference do not need any changes in terms of isotropy or intensity, and can be fed into PlantSeg as it is. Increasing patch size does not improve accuracy. The downside of PlantSeg is that the post-processing algorithms were designed for dense segmentation and therefore tend to over-segment the background, which can be easily fixed by applying a foreground mask or even manually. PlantSeg results in the assignment of very accurate instance masks to most objects, because it finds boundaries of the biological structure of interest and provides a nuclear envelope probability map (Fig. 2D-F). The minor imperfections caused by PlantSeg GASP and final thresholding in PlantSeg segmentation can be very easily improved by removing a few false positives and relabeling a few false negatives.

StarDist-ResNet and StarDist-UNet models output a nuclei probability map (Fig. 2H,K) and nuclei instance segmentation (Fig 2G,I,J,L). Both the StarDist models resulted in very smooth and uniform instance masks of all objects, because it fits star-convex shapes to objects (Fig 2G-L). StarDist is sensitive to object shapes; elongated objects are predicted accurately in its probability maps, but are then sometimes fitted into small and wrong instance masks. The segmentation always looked clean and smooth. Isotropy of data volumes matters, one could specify a grid parameter that downsamples the input to fit instances into the network’s field of view. A bigger patch size can help in terms of object detection but not mean average precision. The imperfection caused by size and shape prior in StarDist segmentation can be improved by merging a few oversegmented instances.

For Cellpose, we fine tuned two pretrained models (Nuclei, Cyto2) and in addition, trained a new model from scratch (Fig. S1). Due to the 2D nature of Cellpose, it is recommended that data for either training or future inference be transformed into isotropic volumes for best results. Cellpose is very sensitive to its diameter parameter. In this study, the fixed default object diameter parameters for pretrained models were set to be 30 for non-nucleus models and 17 for nucleus models, and that for scratch-trained models is inferred from our data. Cellpose results in good instance masks (Fig. S1) but overall less accurate segmentations compared to proposed StarDist and PlantSeg models (Table 2). Overall, while final Cellpose output turned out to be worse than StarDist and PlantSeg even after retraining, it’s important to remember that it was the best method (Table S2) to provide a starting point in absence of human-annotated ground truth in the first step of our experiments.

### Wide applicability of the PlantSeg_3Dnuc and StarDist-ResNet platinum models

So far, the results indicated that PlantSeg_3Dnuc_gold and StarDist-ResNet_3Dnuc_gold emerged as the preferred models for accurately segmenting 3D plant nuclei. Therefore, we trained two final platinum models based on PlantSeg and StarDist-ResNet, respectively, using all available training datasets (Fig. S2). This resulted in the two 3D platinum models, PlantSeg_3Dnuc_platinum and StarDist-ResNet_3Dnuc_platinum. For nuclei segmentation using the two platinum models, we made available the GoNuclear repository (https://github.com/kreshuklab/go-nuclear) that hosts the pipelines used in this study.

To test the broad applicability of the trained platinum models in 3D nuclear segmentation, we used both platinum models to segment nuclei from diverse and challenging datasets, including a variety of tissues from different plant species as well as early mouse embryos, stained with nuclear stains or expressing nuclear reporters. Our diverse 3D nuclei datasets included a fixed, cleared, TO-PRO-3-stained *Antirrhinum majus* ovule; a fixed, cleared, DAPI-stained *Arabidopsis thaliana* ovule; live Arabidopsis sepal nuclei expressing the pATML1::mCitrine-ATML1 reporter (Meyer et al., 2017); live *Cardamine hirsuta* leaf expressing the ChCUC2g::VENUS reporter (Rast-Somssich et al., 2015); and fixed and cleared Arabidopsis shoot apical meristem nuclei expressing the pFD:3xHA-mCHERRY-FD reporter (Cerise et al., 2023; Martignago et al., 2023). In addition, we segmented nuclei of the BlastoSPIM data set obtained by live 3D imaging of blastocyst stage mouse embryos expressing the nuclear marker H2B-miRFP720 using Selective Plane Illumination Microscopy (SPIM) (Nunley et al., 2023).

Both the PlantSeg_3Dnuc_platinum and StarDist-ResNet_3Dnuc_platinum models resulted in comparable high quality segmentations. The results of segmentation using StarDist-ResNet_3Dnuc_platinum are presented here, as its use is less involved compared to PlantSeg_3Dnuc_platinum (Fig. 3, Fig. S3).

**Fig. 3.**
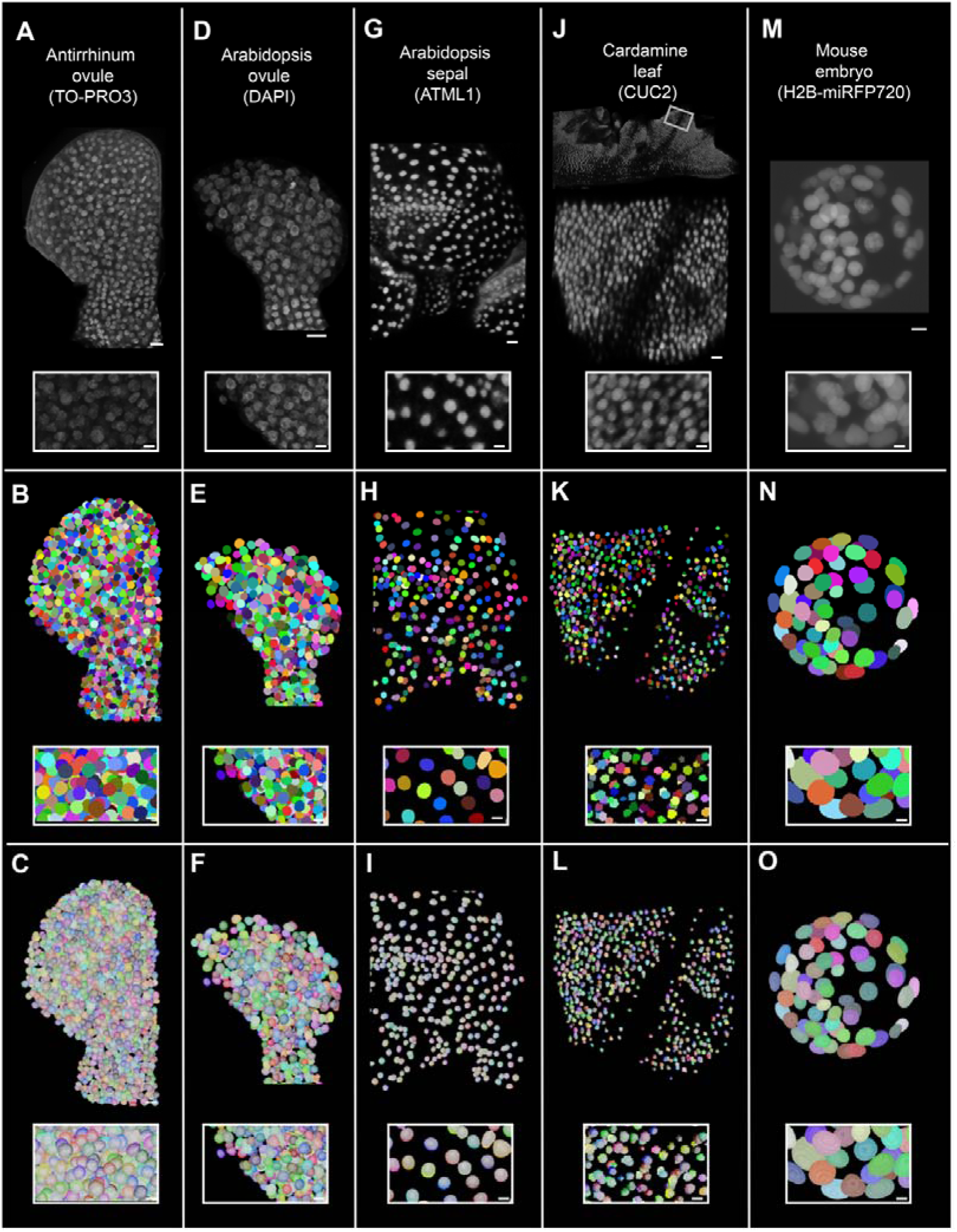
Wide applicability of trained nuclei segmentation models in segmenting stained or nuclear reporter-expressing different plant organ nuclei imaged under different conditions. (A-C) *Antirrhinum majus* ovule nuclei stained with TO-PRO-3 iodide, (D-F) *Arabidopsis thaliana* ovule nuclei stained with DAPI, (G-I) Arabidopsis sepal nuclei expressing the pATML1::mCitrine-ATML1 reporter, (J-L) *Cardamine hirsuta* leaf nuclei expressing the pChCUC2g::Venus reporter, (M-O) Mouse embryo nuclei expressing the H2B-miRFP720 reporter. (A,D,G,J,M) 3D confocal images of raw nuclei stained with a nuclear stain or expressing nuclear reporter. Raw images have been adjusted for brightness and contrast for depiction. (B,E,H,K,N) 3D nuclear segmented stacks, segmented using the StarDist-ResNet model generated from this study. Nuclei IDs are represented in different colors. (C,F,I,L,O) Overlay of 3D segmented stack with the corresponding MorphoGraphX-generated 3D nuclear mesh. (A-O) Insets with white outline show the zoomed-in view of 3D nuclei. Scale Bars: 10 μm (organs) and 5 μm (insets).

We segmented the above-mentioned datasets using the StarDist-ResNet and PlantSeg platinum models after image preprocessing (Table 3). The preprocessing was required to ensure the datasets to be segmented matched the training datasets in nuclear size and quality. We observe that the nuclei of all mentioned datasets could be properly 3D segmented using the proposed models (Fig. 3). Further, even though the models were trained on cleared, high-resolution datasets, they are capable of segmenting nuclei from low resolution datasets as well, for instance the Cardamine leaf nuclei and mouse embryo nuclei from live samples. A precise segmentation of the pChCUC2g::Venus nuclear signal further allows for quantification of the number of pChCUC2g::Venus expressing nuclei along with signal quantification if required. The StarDist-ResNet platinum model could also segment extremely challenging datasets with high variation in intensities after applying some preprocessing (Fig. S3). The results demonstrate the broad applicability of the platinum models in 3D segmentation of nuclei of different tissues and species.

**Table 3.** Datasets used for wide applicability of the proposed method in segmenting different organs and different fluorescent signal types acquired at different resolutions.

| Organism | Organ | Nuclear stain/fluorescent reporter signal | Microscopy | Raw data voxel size (xyz $\mu\text{m}^3$) | Post processing |
| --- | --- | --- | --- | --- | --- |
| *Arabidopsis thaliana* | Shoot apical meristem | pFD:3xHA-mCHERRY-FD | CLSM, 40x 1.25NA Gly objective, cleared sample | 0.242 x 0.242 x 0.4 | Median filtering |
| *Cardamine hirsuta* | Leaf | pChCUC2g::Venus | CLSM, 16x 0.6NA water dipping objective, live sample | 0.498 x 0.498 x 0.5 | Upsampled to 0.125 x 0.125 x 0.25 |
| *Antirrhinum majus* | Ovule | TO-PRO-3 iodide | CLSM, 63x 1.3NA Gly objective, cleared sample | 0.126 x 0.126 x 0.33 | Downsampled to 0.25 x 0.25 x 0.33; smooth 2x |
| *Arabidopsis thaliana* | Ovule | DAPI | CLSM, 63x 1.3NA Gly objective, cleared sample | 0.063 x 0.063 x 0.27 | Downsampled to 0.25 x 0.25 x 0.28; smooth 2x |
| *Arabidopsis thaliana* | Sepal | pATML1::mCitrine-ATML1 | CLSM, 20x 1.0NA Water objective | 0.276 x 0.276 x 0.8 | Autobright; smooth 3x |
| *Mouse* | Early embryo | H2B-miRFP720 | SPIM | 0.208 x 0.208 x 2 | Downsampled in x, y and unchanged in z to 0.6 x 0.6 x 2 |

The table summarizes the organism, organ, type of signal, microscopic method, image voxel size and any preprocessing applied to optimize the segmentation of the wide applicability dataset.

### MorphoGraphX as a platform for mapping 3D nuclei to whole organ cell atlas with single cell and tissue resolution

Multichannel 3D confocal imaging allowed simultaneous imaging of both the cell and nuclear stain channels. MorphoGraphX enables 3D visualization and allows complex annotations and quantifications (Fig. S2F-I, Fig. S3H). We reasoned that it should be possible to combine 3D cell segmentation and 3D nuclear segmentation of the imaged 3D stack. 3D cell segmentation assigns cells their cell IDs and 3D nuclear segmentation assigns nuclei their nuclei IDs; however, they are not directly linked. In MorphoGraphX, these 3D cell and nuclei segmentation images are converted to 3D meshes representing individual objects. To address the issue of linking nuclei and corresponding cell IDs, we developed a novel process in MorphoGraphX that automatically annotates and links nuclei IDs with their corresponding cell IDs (Fig. 4A-H) (see Materials and Methods). The 3D cell meshes can then be assigned tissue labels via manual or semi-automated cell-type labeling (Strauss et al., 2022) (Fig. 4A-B).

**Fig. 4.**
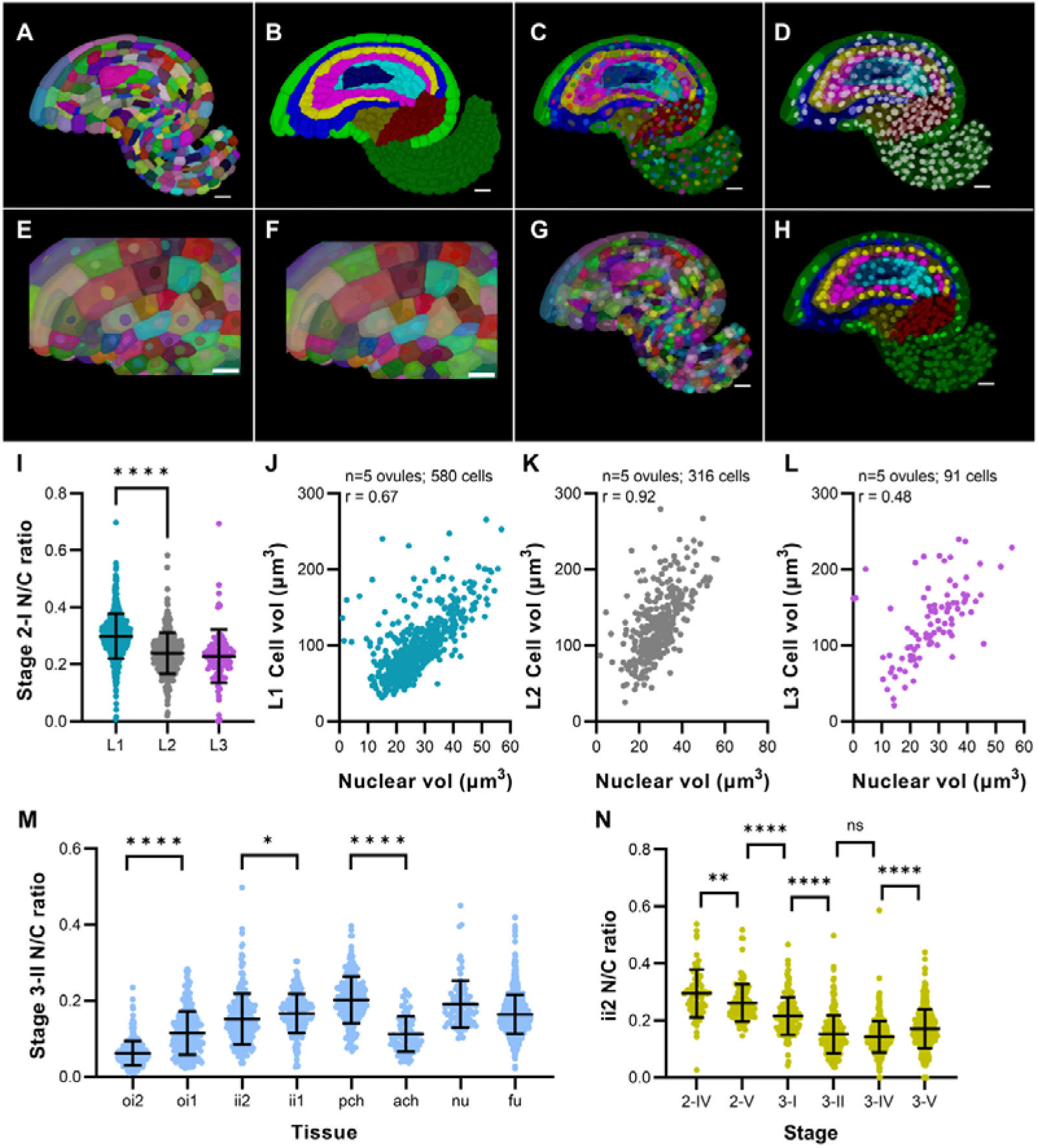
MorphoGraphX as a platform for mapping 3D nuclei to whole organ cell atlas at single cell and tissue resolution. (A-H) Stage 3-II 3D cell and nuclei meshes for the same ovule sample generated from corresponding segmented stacks. (A) Mid-sagittal section of 3D mesh showing cell IDs in different colors. (B) Mid-sagittal section of 3D mesh showing cell parent (tissue) labels. (C) Cell-type labeled 3D mesh overlaid with nuclei mesh showing nuclei IDs in different colors. (D) Cell-type labeled 3D mesh overlaid with nuclei mesh showing nuclei lacking parent labels. (E) Cropped section of 3D mesh showing that initially cell IDs are initially independent of nuclei IDs (cells and their corresponding nuclei in different colors). (F) Cropped section and (G) mid-sagittal section of 3D mesh showing cell IDs mapped onto their corresponding nuclei using MorphoGraphX processes, resulting in the same color for cells and their corresponding nuclei. (H) In the final step parent tissue labels of cells are mapped onto the corresponding nuclei in MorphoGraphX. (I) Plot showing N/C ratio of the radial layers, L1, L2, and L3 of stage 2-I ovule primordia. (J-L) Plots showing correlation between nuclear and cell volumes in different layers of stage 2-I primordia along with the respective Pearson correlation coefficients, *r*. (J) L1, (K) L2, (L) L3. (M) Plot showing nuclear to cell volume ratio (N/C) of different tissues and tissue layers of stage 3-II ovules. (N) Plot showing N/C ratio of the outer layer of the inner integument (ii2) for different stages of ovule development from 2-IV up to 3-V. Asterisks represent statistical significance (ns, p≥0.5; *, p<0.05; **, p<0.01, ***, p<0.001; ****, p<0.0001; Student’s t-test). Scale bars: 10 μm.

In addition to linking nuclear and cell IDs, we also added MorphoGraphX tools to quantify the Euclidean distance between 3D cell and 3D nuclear centroids or map 2.5D cells to underlying 3D nuclei (Fig. S4) (Materials and Methods). 3D cell segmentation can be challenging especially when working with live images. In such cases, one may have to resort to 2.5D cell segmentation. We present a MorphoGraphX method for associating 2.5D surface cells with 3D nuclei.

MorphoGraphX achieves this link by projecting 3D segmented nuclei stacks onto the 2.5D segmented cell mesh (Fig. S4A-C). Additionally, the process “Select Duplicated Nuclei” is a useful tool to identify cell segmentation errors as it detects cells with more than one nucleus. This entire collection of processes are included in MorphoGraphX version 2.0.2. and higher and can be found in the process folder “Mesh/Nucleus” (see Materials and Methods). The development of these new MorphoGraphX processes opens up new possibilities to integrate cell features with nuclei features and to study quantitative parameters of nuclei in their cellular context.

### Developmental regulation of the nucleus-to-cell volume ratio in Arabidopsis ovules

For more than a century it has been noticed that the N/C ratio is a constant parameter of a given cell type that can vary between cell types in multicellular organisms (Cantwell and Nurse, 2019c; Wilson, 1925). Most of these studies involved selecting a few cells of embryos or single cells, such as yeast, and measurements based on diameter or area values derived from 2D sections. Here, we investigated the N/C ratio in Arabidopsis ovules of different stages and in full 3D tissue context. We measured the nuclear volumes, cell volumes, N/C ratios and their trends in five stage 2-I ovule primordia and in two more differentiated stage 3-II ovules. In addition, we assessed these parameters during the development of an integumentary cell layer using two ovules per stage (Fig. 4I-N) (Fig. S5).

The dome-shaped Arabidopsis ovule primordium, like the shoot apical meristem, has a layered organization, such that the L1, L2, and L3 are the outer to inner layers, respectively (Jenik and Irish, 2000; Satina et al., 1940; Schneitz et al., 1995). At stage 2-I the primordium is further characterized by the presence of an enlarged L2-derived megaspore mother cell (MMC) at the tip that will undergo meiosis and eventually produce the haploid female gametophyte (Schneitz et al., 1995; Vijayan et al., 2021). We investigated if nuclear and cell volumes, as well as N/C ratios differ in a layer-specific manner in the ovule primordium. We observed that the L1 layer can be distinguished from the L2 and L3 layers by its different N/C ratio, as the N/C ratio of L1 cells was statistically different from the N/C ratio of L2 or L3 cells. The L2 and L3 N/C ratios were not noticeably different (Fig. 4I, Fig. S5A,B). Cells of the outermost L1 layer have the highest N/C ratio (0.30 ± 0.08 (mean ± SD), followed by the cells of the inner L2 (0.24 ± 0.07) and L3 (0.23 ± 0.09) layers (Fig. 4I). For all three layers, we obtained a positive Pearson correlation coefficient, r, between nuclear and cell volumes; the correlation is strongest in the L2 layer, followed by the L1 and L3 layers, respectively (Fig. 4J-L). When analyzing the average cell and nuclear volumes for each layer, we found that the average cell volumes of the L2 (128.10 ± 47.73 µm^3^, excluding MMCs) and L3 (132.70 ± 54.53 µm^3^) layers were similar and markedly larger than the average cell volume of the L1 (98.86 ± 38.06 µm^3^) layer (Fig. S5A). In contrast, the average nuclear volume between the three cell layers remained comparable with values of 27.88 ± 9.14 µm^3^ (L1), 28.92 ± 8.87 µm^3^ (L2, excluding MMCs), and 27.44 ± 10.53 µm^3^ (L3) (Fig. S5B). Thus, the difference in the N/C values between the L1 and L2/L3 layers relates to the smaller average cell volume in the L1 compared to the L2 and L3 layers.

Current evidence suggests that nuclear size scales with cell size and not with the amount of nuclear DNA (Cantwell and Nurse, 2019c). We tested if the scaling rule holds true for the MMCs (Fig. S5C,D). We found that the average nuclear and cell volumes of the tested MMCs (147.9 ± 27.85 µm^3^ for the nuclear volume and 845.4 ± 101.5 µm^3^ for the cell volume) both exceeded the respective values of the other much smaller L2 cells by approximately a factor of 5. As a result, the N/C ratio values of the MMCs and the other L2 cells were indistinguishable, and thus the MMCs conform to this rule.

To confirm the finding of cell type-specific N/C ratios in the ovule primordium we explored the more differentiated stage 3-II ovules exhibiting a clear multi-tissue organization. By this stage the Arabidopsis ovule is composed of the distal nucellus, which contains the developing female gametophyte, the central chalaza with two lateral determinate structures, the integuments, and the proximal funiculus, the stalk that connects the ovule to the placenta (Schneitz et al., 1995; Vijayan et al., 2021). In addition, the chalaza can be divided into an anterior and posterior chalaza based on morphological criteria such as different cell shapes and sizes of its constituent cells. In addition, each integument consists of two cell layers, each one cell thick. The analysis of the average N/C values across different tissues revealed that the nucellus and funiculus exhibited comparable values. In contrast, we found the posterior chalaza to show a higher N/C ratio than the anterior chalaza (Fig. 4M). We also observed that the inner layers of both the outer and inner integuments exhibited a higher N/C ratio than the corresponding outer layers.

To address the question if the N/C ratio changes during development of a specific tissue layer, we focussed on the outer layer of the inner integument (ii2). We analyzed the ii2 nuclear and cell volumes, and the N/C ratios for stages 2-IV, 2-V, 3-I, 3-II, 3-IV and 3-V. We observed that from stage 2-IV to stage 3-IV, there was a decline in the ii2 N/C ratio (0.29 ± 0.08 towards 0.15 ± 0.07), followed by an increase from stage 3-IV to 3-V (0.14 ± 0.06 versus 0.17 ± 0.07) (Fig. 4N). To assess the basis for this decrease in the N/C ratio during development of the ii2 layer we analyzed the average nuclear and cell volumes between successive stages (Fig. S5C-D). We found that the average cell volume of ii2 cells increased noticeably with a value of 129.8 ± 58.06 µm^3^ at stage 2-IV and 220.50 ± 130.0 µm^3^ at stage 3-IV (Fig. S5C). In comparison, the average nuclear volume experienced only minor alterations (35.46 ± 13.73 µm^3^, stage 2-IV; 27.03 ± 10.36 µm^3^, stage 3-II; 31.93 ± 9.91 µm^3^, stage 3-V) (Fig. S5C). Thus, we find that the change in the N/C ratio during development of the ii2 cell layer is related to a marked increase in cell volume accompanied by a largely constant nuclear volume. Further estimation of the stagewise Pearson correlation coefficient, *r*, for ii2 revealed that there is a positive correlation between cell volumes and corresponding nuclear volumes of ii2 across development up to stage 3-IV. By stage 3-V, however, this correlation is noticeably reduced (Fig. S5E-J).

In summary, the results suggest that the N/C ratio is specific to a cell type and its developmental stage in the Arabidopsis ovule.

### Automatic proofreading of 3D cell segmentation based on reliable 3D nuclear segmentation

Despite the significant improvement in cell boundary prediction provided by the PlantSeg segmentation pipeline, the final image segmentation may still contain some errors in certain regions of the images where cell wall staining is poor. An example is the faint walls around the megaspore mother cell (MMC) in young Arabidopsis ovules (Vijayan et al., 2021) (Fig. 5A-D). From the raw cell wall stain images, it is almost impossible to identify the presence of these faint walls. A similar scenario sometimes applies to cells in the interior chalaza (Fig. 5E-H). The processed raw images (brightened) along with the nuclei stain clearly display the faint wall and the presence of multiple nuclei in this region confirming the cell segmentation error in this region.

**Fig. 5.**
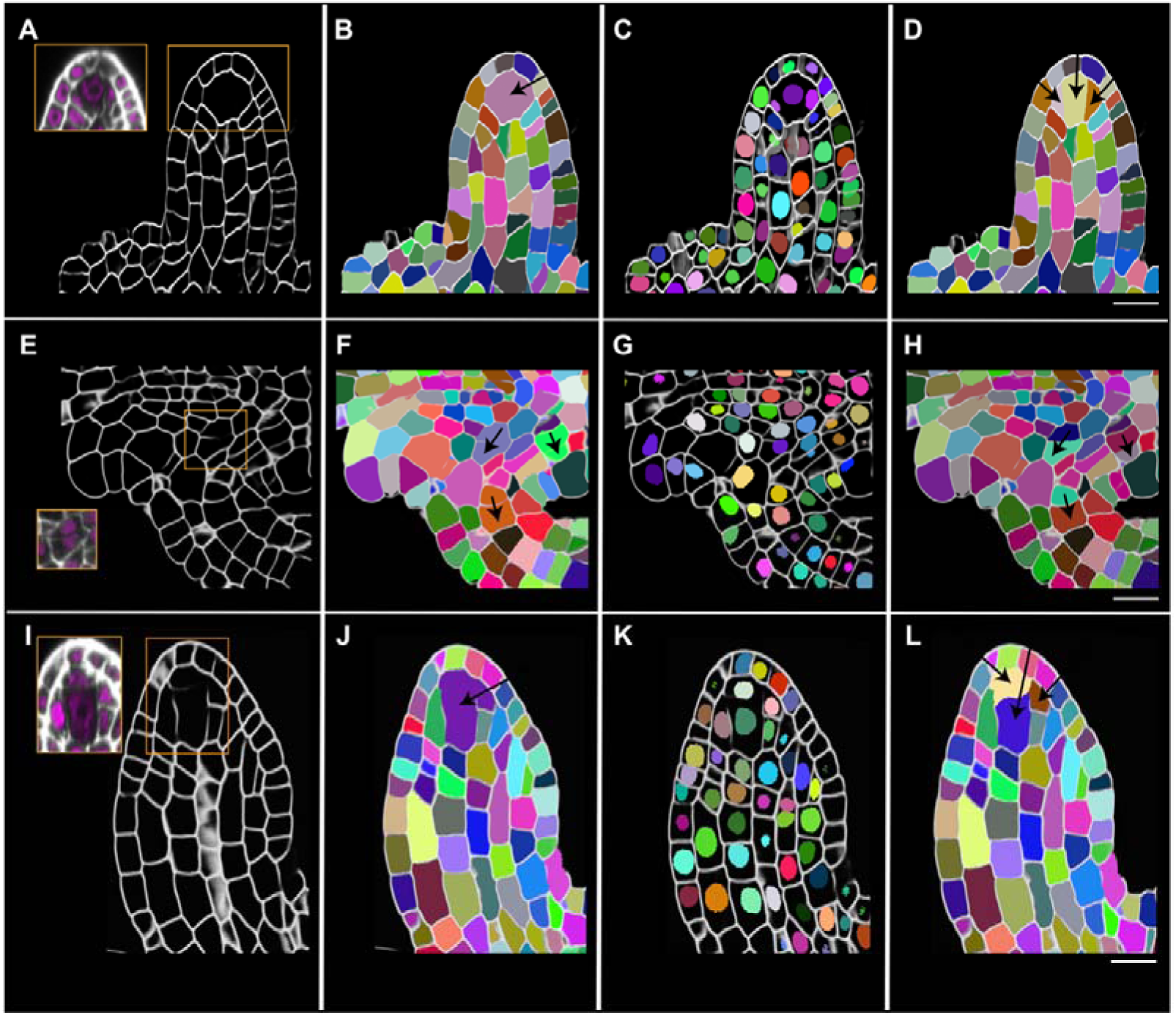
PlantSeg proofreading tools to correct 3D cell segmentation errors. (A-D) Mid-sagittal section *Arabidopsis thaliana* ovule primordium (dataset 598A, (Vijayan et al., 2021)). (E-H) Cropped section of an *Arabidopsis thaliana* 3-II ovule (dataset 527, (Vijayan et al., 2021)). (I-L) Mid-sagittal section of a *Cardamine parviflora* ovule primordium (dataset 1598B, Mody et al., 2023). (A,E,I) 3D cell boundary predictions along with insets showing raw SR2200 (white) and TO-PRO-3 channel (magenta) signals after adjusting for brightness and contrast to show the weak cell wall staining in specific regions (outlined in orange boxes) and resulting in missing or incomplete walls in the cell boundary predictions. (B,F,J) Plant-seg cell segmentations overlaid with cell boundary prediction. Black arrows point to undersegmented cells. (C,G,K) StarDist-segmented nuclei overlaid with cell boundary prediction, showing multiple nuclei in the undersegmented cells in the MMC region (B,J) and in cells of funiculus and chalaza (F). (D,H,L) 3D cell segmentations corrected with PlantSeg proofreading tools (black arrows) and overlaid with the cell boundary prediction. *Cardamine parviflora* ovule primordia are crassinucellate (K,L); the ability to visualize this is lost after cell segmentation (I,J). PlantSeg proofreading tools enable re-distinguishing the primary parietal cell from the MMC. Scale Bars: 10 μm.

We developed a python script called “proofreading” to automatically correct the instance 3D cell segmentation using a trusted and proofread 3D nuclear segmentation and added it to the growing collection of helper tools for the PlantSeg pipeline (https://github.com/hci-unihd/plant-seg-tools). The script takes the cell boundary prediction, cell segmentation and nuclei segmentation as input images. It automatically finds the erroneous cell segmentation by first quantifying the number of nuclei within a cell. When it finds a cell with more than one nucleus, a bounding box is approximated in 3D around this cell. Further corrections are only made within the bounding box. Corrections are made by resegmenting the erroneous 3D cell using watershed segmentation with nuclei as seeds. The t-merge parameter can be altered to improve the segmentation further if the default value does not seem to improve the result. The method does not apply to a scenario where the segmentation error relates to a missing cell instead of an under segmented cell. The detailed method is described in the Materials and Methods section.

This method now corrects the segmentation error in most cases and leaves other cells without segmentation errors untouched (Fig. 5C,D and G,H). We performed another test by assessing *Cardamine parviflora* (*C. parviflora*) ovule primordia (Fig. 5I-L).

This species harbors weakly crassinucellate ovule primordia (Endress, 2011), i.e., it develops an additional hypodermal cell layer, with an initial archesporial cell in the L2 undergoing periclinal division resulting in an upper parietal cell and a lower MMC (Harvey and Smith, 2013; Mody et al., 2023). The ability to visualize this is usually lost after standard PlantSeg-based 3D cell segmentation (Fig. 5I,J), but the proofreading script can correct this error (Fig. 5K,L). The proofreading thus minimizes 3D cell segmentation errors and enables the examination of 3D cell volumes for cells that are challenging to segment accurately.

## Discussion

We present a collection of computational tools and datasets that extend the capabilities for quantitative analysis of 3D digital organs. We have developed a deep-learning based computational toolkit for 3D nuclear segmentation that enables accurate 3D segmentation of nuclei in a variety of 3D digital organs labeled with a range of nuclear markers or stains, even in faintly stained and noisy images.

Importantly, we not only provide a valuable plant nuclear dataset for training 3D nuclear segmentation algorithms but also two accurate platinum models for 3D nuclear segmentation with broad applicability. In addition, we outline novel and processes that we have added to MorphoGraphX to enable the analysis of various cell-nucleus geometric parameters in 3D, including the N/C ratio. Finally, we have created a proofreading script that significantly improves the fidelity of 3D cell segmentation. All tools are open source and readily available to the community via public software repositories.

A particular value of the 3D nuclear segmentation toolkit lies in its broad applicability. The method can be successfully used with various nuclear staining methods, ranging from different nuclear stains with variable staining intensities, such as TO-PRO-3 or DAPI, to nuclear reporters based on fluorescent reporters. In addition, nuclei can be segmented in data sets obtained from cleared or live tissue, not only from a range of different plant tissues, but also from animal tissues such as mouse embryos. An optimized workflow from imaging to 3D segmentation of nuclei dataset can be found in the Materials and Methods section.

We used PlantSeg (Wolny et al., 2020), Cellpose (Stringer et al., 2021) and StarDist (Schmidt et al., 2018; Weigert et al., 2020) as three strong baselines for 3D nuclear segmentation and performed a comparative analysis of the performance of the models obtained from each platform. Cellpose was the only tool that provided a pre-trained model which could perform the initial segmentation. However, in the presence of ground truth, it was demonstrated to be less stable, with more variability in the results depending on the training/test split of the data. Re-trained PlantSeg and StarDist both demonstrated excellent, stable performance. The advantage of PlantSeg is its ability to also perform cell segmentation from membrane staining and the general absence of explicit star-convexity prior which can be harmful for segmentation of irregular nuclei. However, in very noisy conditions StarDist is preferred as the shape prior helps it overcome the low SNR. It also needs to be noted that our ground truth annotations are produced through iterative improvement using a StarDist model, so the resulting shapes might be biased towards being more regular and star-convex.

An important feature of MorphoGraphX is the projection of secondary signals onto the cell surfaces, which enables the quantification of nuclei, cell wall or cytoplasmic signal intensities based on the cellular segmentation (Barbier de Reuille et al., 2015; Montenegro-Johnson et al., 2015). What up to now was missing, however, was the integration of the size and shape of the nuclei into the cellular framework. We present an extension to MorphoGraphX that includes the ability to assign individual nuclei to their corresponding 3D cells in digital 3D organs. To this end, we have developed a number of processes that are part of the latest versions of MorphoGraphX from version 2.0.2. This improvement allows the analysis of various relationships between the nucleus and the cell it is located in, in 3D, including determination of the Euclidean distance between cell and nuclear centroids, mapping 2.5D cells to underlying 3D nuclei, or identification of more than one nucleus in a cell. For example, quantification of the Euclidean distance between cell centroids and nuclear centroids was instrumental in developing the notion that positioning the plane of cell division in cells of the early Arabidopsis embryo does not depend on the precise position of the nucleus (Vaddepalli et al., 2021).

The “Kernplasma-Relation” (nucleus-cytoplasm relation) has fascinated cell biologists since its discovery around the turn of the last century (Conklin, 1912; Hertwig, 1903; Strasburger, 1893; Wilson, 1925). The currently favored model states that nuclear size scales with cell size and that the N/C ratio is cell-type specific (Cantwell and Nurse, 2019c). Our findings in the Arabidopsis ovule support this notion. For example, we noticed that the outermost L1 layer has a larger N/C ratio compared to the L2 and L3 layers in the ovule primordium. This change is largely due to alterations in cell not nuclear size. Thus, we find that similarly sized nuclei can populate cells with significant size differences, supporting the notion that this scaling rule is valid in the context of a specific cell type. Interestingly, this result differs from the scenario in the L1, L2, and L3 layers of the Arabidopsis shoot apical meristem (SAM), where cells of the three layers have similar N/C ratios (Wenzl and Lohmann, 2023), further highlighting the tissue specificity of N/C ratios. The observed changes in the N/C ratio during ii2 development may indicate early changes in the differentiation status. For example, threshold values of N/C ratios in Xenopus oocytes have been shown to be critical for transcriptional initiation associated with developmental stage transition (Jevtić and Levy, 2015).

How nuclear size is regulated is poorly understood (Cantwell and Nurse, 2019c). Current evidence indicates that nuclear size in yeast is controlled by several processes, including osmotic forces, bulk nucleocytoplasmic transport, transcription and RNA processing, linker of nucleoskeleton and cytoskeleton (LINC) complexes, and membrane expansion (Cantwell and Nurse, 2019a; Cantwell and Nurse, 2019b; Deviri and Safran, 2022; Lemière et al., 2022). In Arabidopsis, two nuclear envelope proteins were described to function redundantly in the control of nuclear size and shape in response to hyperosmotic stress in root tip cells (Goswami et al., 2020). The straightforward tools presented for the quantitative study of nuclear volume will facilitate the functional dissection of the control of nuclear size and shape in multicellular organisms such as seed plants.

Finally, the PlantSeg-based cell segmentation proofreading script provides a useful tool to correct 3D cell segmentation errors due to weak cell wall staining. The method uses the successfully 3D segmented nuclei as seeds and thus its success critically depends on precise 3D nuclear segmentation. Our results indicate that it can dramatically improve the fidelity of 3D cell segmentation, as indicated by the observed corrections of the notoriously difficult to segment cells surrounding the MMC in *A. thaliana* and *C. parviflora* ovule primordia.

In conclusion, the novel computational toolkit we present here augments the growing suite of tools that enable the generation and detailed quantitative analysis of 3D digital organs at single cell resolution.

## Materials and Methods

### Plant work and transformation

*Arabidopsis thaliana* (L.) Heynh. var. Columbia (Col-0), *Cardamine parviflora,* and *Antirrhinum majus* were used as the wild-type strains. Plants were grown as previously described (Fulton et al., 2009). Arabidopsis Col-0 plants were transformed with the pUBQ::H2B:tdTomato construct using Agrobacterium strain GV3101/pMP90 (Koncz and Schell, 1986) and the floral dip method (Clough and Bent, 1998). Transgenic T1 plants were selected on Hygromycin (20 mg/ml) or Sulfadiazine (5 µg/ml) plates according to the selection.

### Recombinant DNA work

For DNA work, standard molecular biology techniques were used. PCR fragments used for cloning were obtained using Q5 high-fidelity DNA polymerase (New England Biolabs, Frankfurt, Germany). All PCR-based constructs were sequenced. Constructs were generated using the GreenGate system (Lampropoulos et al., 2013). pUBQ::H2B:tdTomato: a dual reporter for cell membrane and H2B nuclei was designed and constructed using GreenGate. pUBQ::H2B:tdTomato and pSUB::gSUB:mTurquoise2 were assembled into the intermediate vectors and then combined into the pGGZ0001 destination vector with a standard GreenGate reaction. The pSUB::gSUB:mTurquoise2 expression was weak or absent and we only imaged H2B nuclei in this study. Half MS plate containing Sulfadiazine (5 µg/ml) was used for plant resistance selection.

### Clearing and staining of ovules

Fixing, clearing, and staining of dissected ovules was performed as described earlier (Tofanelli et al., 2019).

### Microscopy and data acquisition

Confocal laser scanning microscopy of ovules of *Arabidopsis thaliana*, *Cardamine parviflora*, and *Antirrhinum majus* stained with SR2200 and TO-PRO-3 iodide or DAPI was performed on an upright Leica TCS SP8 X WLL2 HyVolution 2 (Leica Microsystems) equipped with GaAsP (HyD) detectors and a 63x glycerol objective (HC PL APO CS2 63×/1.30 GLYC, CORR CS2). Laser power or gain was adjusted for z compensation to obtain an optimal z-stack. SR2200 fluorescence was excited with a 405 nm diode laser (50 mW) with a laser power ranging from 0.1 to 1.5% intensity and detected at 420 to 500 nm with the gain of the HyD detector set to 20. TO-PRO-3 iodide fluorescence excitation was done at 642 nm with the white-light laser with a laser power ranging from 2 to 3.5% and detected at 655 to 720 nm with the gain of the HyD detector set to 200. For z-stacks 8, 12 or 16-bit images were captured at a slice interval of 0.28 μm or 0.33 μm with optimized system resolution of 0.126 μm × 0.126 μm × c μm (c=0.280 or 0.330) as final pixel size according to the Nyquist criterion. Scan speed was set to 400 Hz, the pinhole was set to 0.6 to 1.0 Airy units, line average was between 2 and 4, and the digital zoom was set between 0.75 and 2, as required. Laser power or gain was adjusted for z compensation to obtain an optimal z-stack. Image acquisition parameters for the pUBQ::H2B:tdTomato reporter line: SR2200; 405 diode laser 0.10%, HyD 420–480 nm, detector gain 10. tdTomato; 554 White laser 4%, HyD 570–630 nm, detector gain 80. TO-PRO-3; 642 nm White Laser 2%, HyD 660–720 nm, detector gain 100. In each case sequential scanning was performed to avoid crosstalk between the spectra. DAPI stained ovules were excited with a 405 diode laser 3 %, HyD 420–480 nm, detector gain 100.

Confocal images of live *Cardamine hirsuta* Oxford leaf were performed on an upright Leica TCS SP8 equipped with a 16x 0.6NA multi immersion objective (HC FLUOTAR L 16x/0,60 IMM CORR VISIR). Sample was mounted on a glass slide under a coverslip, stained with 1% Propidium iodide in water for cell wall fluorescence along with ChCUC2g::Venus signal. Venus was excited using a 514 diode laser 2.5%, detected using the HyD 520-560, detector gain 100.

The dataset of pATML1::mCitrine-ATML1 expressing nuclei in the Arabidopsis flower (pATML1mCitrine-ATML1_flower1_t08.tif) has been obtained from (Meyer et al., 2017). The dataset of Arabidopsis shoot apical meristem nuclei expressing the pFD:3xHA-mCHERRY-FD reporter(Cerise et al., 2023; Martignago et al., 2023). The dataset of mouse embryo nuclei (F49_149) has been obtained from (Nunley et al., 2023). 2D, 3D or 2.5D rendered snapshots were taken using MorphoGraphX. Images were adjusted for color and contrast using Adobe Photoshop (Adobe, San José, USA) or MorphoGraphX software (https://www.morphographx.org) (Barbier de Reuille et al., 2015; Strauss et al., 2022).

### Model training and score quantification

The new training dataset (N1-N5) is composed of three image channels: SR2200 cell wall stain, H2B:tdTomato nuclear reporter, and TO-PRO-3 nuclear stain. The SR2200 cell wall stain was processed with the PlantSeg pipeline to generate a 3D cell boundary prediction and segmentation. 3D segmentation of the strong tdTomato nuclei reporter signal was performed using the default Cellpose nuclei model. It was then proofread and used as the “initial ground truth”. This study provides five initial ground truth segmentation datasets (Table 4) for model training. Initial model training was performed using the initial ground truths and trained on the weak TO-PRO-3 channel. The StarDist-ResNet_3Dnuc_initial model was then used to segment the original weak TO-PRO-3-based nuclear stain channel resulting in a modified ground truth which was then human proofread, resulting in the “gold ground truth”. Gold model training was performed using the gold ground truths and trained on the weak TO-PRO-3 channel.

**Table 4.**
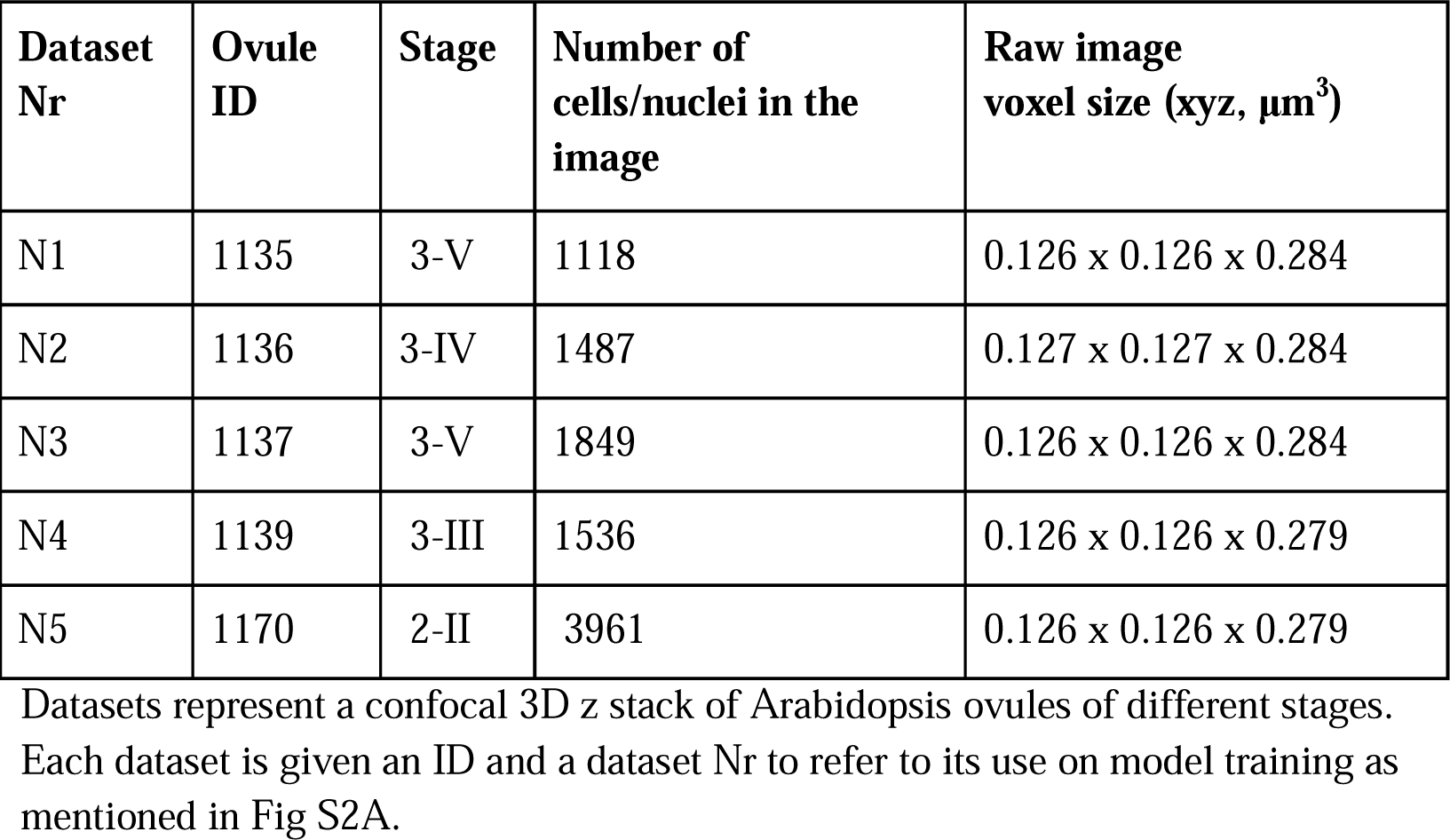
Training and testing dataset for five fold training and for the platinum trained model.

| Dataset Nr | Ovule ID | Stage | Number of cells/nuclei in the image | Raw image voxel size (xyz, $\mu\text{m}^3$) |
| --- | --- | --- | --- | --- |
| N1 | 1135 | 3-V | 1118 | 0.126 x 0.126 x 0.284 |
| N2 | 1136 | 3-IV | 1487 | 0.127 x 0.127 x 0.284 |
| N3 | 1137 | 3-V | 1849 | 0.126 x 0.126 x 0.284 |
| N4 | 1139 | 3-III | 1536 | 0.126 x 0.126 x 0.279 |
| N5 | 1170 | 2-II | 3961 | 0.126 x 0.126 x 0.279 |

For quantitative evaluation of the models, we trained five different models during both “initial” and “gold” training of each of the PlantSeg, StarDist, and Cellpose neural networks. Cross-validation with one datasets kept out for testing was used (Fig S2A), i.e. for model 1, N1-N4 data was used for model training while N5 was the testing dataset. Each model training and testing involved three training datasets, one validation dataset, and one testing dataset. For example, one PlantSeg model was trained on N1, N2, N3 datasets, validated on N4 dataset, and tested on N5 dataset; the next was trained on N2, N3, N4 datasets, validated on N5 dataset, and tested on N1 dataset and so on. Therefore, the trained models from the initial and gold training include 15 (3 X 5) initial models and 30 (6 X 5) gold models (Table 5).

**Table 5.** List of all models.

| Sl | Model name | Tool used | Model | Version | Number of trained models |
| --- | --- | --- | --- | --- | --- |
| 1 | PlantSeg_3Dnuc_initial | PlantSeg | UNet GASP | Initial | 5 |
| 2 | StarDist-ResNet_3Dnuc_initial | StarDist | ResNet | Initial | 5 |
| 3 | Cellpose-Finetune-nuclei_3Dnuc_initial | Cellpose | Finetune nuclei | initial | 5 |
| 4 | PlantSeg_3Dnuc_gold | PlantSeg | UNet GASP | gold | 5 |
| 5 | StarDist-ResNet_3Dnuc_gold | StarDist | ResNet | gold | 5 |
| 6 | StarDist-UNet_3Dnuc_gold | StarDist | Unet | gold | 5 |
| 7 | Cellpose-Finetune-nuclei_3Dnuc_gold | Cellpose | Finetune nuclei | gold | 5 |
| 8 | Cellpose-Cyto2_3Dnuc_gold | Cellpose | Finetune cyto2 | gold | 5 |
| 9 | Cellpose-Scratch_3Dnuc_gold | Cellpose | Train from scratch | gold | 5 |
| 10 | PlantSeg_3Dnuc_platinum | PlantSeg | UNet GASP | Platinum | 1 |
| 11 | StarDist-ResNet_3Dnuc_platinum | StarDist | ResNet | Platinum | 1 |

To evaluate and compare models and settings, mean Average Precision was chosen for scoring (Caicedo et al., 2019). To make clear the exact metric used among many variants (Hirling et al., 2023), the code for evaluation is publicly available to complement the following formulae. Intersection over Union (IoU), or the Jaccard index, measures the overlap between a predicted mask and a ground-truth mask for the testing dataset. It is represented on a scale from 0 to 1, where a value of 1 signifies a perfect match at the pixel level, and a value of 0.5 indicates that the number of correctly matched pixels is equal to the combined number of missed and false positive pixels. We define the precision of the segmentation for an image as *precision(t)* = 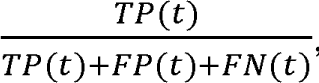, where t is the threshold, TP the number of objects that match true objects with IoU value above t, FP the number of objects that have no true object associated with, and FN the number of true objects that are not present in the segmentation. The average precision (AP) over a range of IoU is defined as 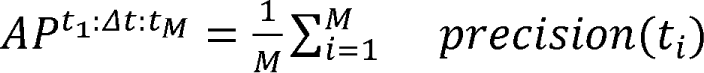 where M is the number of IoU thresholds that i=1 i range from t_1_ to t_M_ with a step size of Lit. As a break from tradition, for each setting, five models were evaluated, each with one image, and the scores were averaged, thus the mean AP in our study is 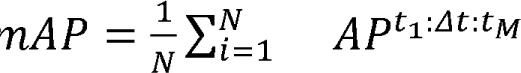, where N is the number of i=1 images and models. A five-fold average precision at 50% IoU across 5 models, denoted as mAP^SO^, is used as a detection score, and a five-fold average precision over {50%, 55%, …, 95%} IoU and across 5 models, denoted as mAP^SO:S:9S^ or simply mAP, is used as the instance segmentation score. The initial and gold models have been quantified using the AP scores and reported along with standard deviation. The initial models trained on PlantSeg, StarDist-ResNet, and Cellpose-Finetune-Nuclei and the gold models trained on PlantSeg, StarDist-ResNet, StarDist-UNet, Cellpose-Finetune-Cyto2, Cellpose-Finetune-Nuclei, and Cellpose trained from scratch were evaluated with five-fold AP scoring (Tables 1 and 2). Detailed quantification of AP scores for evaluation of segmentation can be found in the Supplementary File 1.

Finally, two robust and widely applicable platinum models are proposed where all five datasets (N1-N5) were used for training final robust models: PlantSeg_3Dnuc_platinum and StarDist-ResNet_3Dnuc_platinum (Fig S2, Fig 3, Fig 4). We provide the two platinum models through the BioImage Model Zoo for FAIR use through different client tools of our community. For the sake of reproducibility, we also provide the full bundle of models we trained: initial, gold and platinum, to be downloaded from Biostudies repository S-BIAD1026 (Table 5).

### Data preprocessing for performing segmentation using the proposed StarDist, Cellpose and PlantSeg models

For the best performance, StarDist requires the raw data to be rescaled so that the median diameter of nuclei fits into the field of view of the model. We recommend resampling the dataset to a voxel size of 0.25 x 0.25 x 0.25 µm³ (xyz) for the StarDist-ResNet platinum model proposed by this study. The grid parameter in the config is a StarDist model parameter that specifies the downsampling factor in each dimension; [2, 4, 4] downsamples the image by 2 in z and by 4 in x and y. Cellpose models need to know the diameter or an estimate of that to match the testing datasets’ objects to original datasets’ object diameter (30 for cell models and 17 for the nuclei model); PlantSeg model does not require rescaling to match object size, but it is recommended to match the voxel size to 0.25x 0.25 x 0.25 µm³ (xyz) so that the membrane has similar thickness. This paper comes with data, code, models and configuration files.

### Mapping cell labels to nuclei labels in MorphoGraphX

3D cell and nuclei meshes were generated from segmented stacks using the Marching cubes 3D process with a cube size of 0.5 μm for fine details. Cell-type labeling assigns parent (tissue) labels to the cell IDs. Cell-type labeling was done as described in (Vijayan et al., 2021). The cell and nuclear volumes were obtained using the “Mesh/Heat Map/Analysis/Cell Analysis 3D” process in MorphoGraphX. Initially, the cells in the 3D cell mesh (Mesh 1) have their unique cell IDs and the nuclei in the 3D nuclear mesh (Mesh 2) have their unique nuclei IDs. Both the IDs are mapped using the MorphoGraphX process “Mesh/Nucleus/ Label Nuclei”. In detail, this process identifies the cells in which nuclei are located. It is run on the active 3D cell mesh in MorphoGraphX mesh 1, while the 3D nuclei mesh is loaded in the MorphoGraphX Mesh 2. The process assigns cell IDs as “parents” annotation to the nuclei labels, thereby linking cells IDs to nuclei IDs. On the 3D nuclei mesh (active), the “Mesh/Lineage tracking/Save parents” process was used to save the nuclei IDs and their corresponding parent cell IDs in a csv file, followed by the “Mesh/Lineage tracking/Copy parents to labels” process to rewrite the nuclei labels IDs to that of cells. These processes in combination with “Mesh/Heat map” and “Mesh/Heat map/Operators/Export heat to Attr Map” processes were used to generate csv files containing cell IDs, their corresponding nuclei IDs, parent (tissue) labels, and cell and nuclei geometric attributes.

Further, we created a process (“Mesh/Nucleus/Select Duplicated Nuclei”) to detect and automatically select nuclei in cells where multiple nuclei were detected. This process was used to detect segmentation errors. Another process (”Mesh/Nucleus/Distance Nuclei”) was implemented to quantify the Euclidean distance between cell centroids and nuclei centroids. We also included a process (“Mesh/Nucleus/Label Nuclei Surface”) to associate 3D segmented nuclei IDs with the cells of curved surface meshes. All these processes are documented within MorphoGraphX (Help/Process Docs). Specific application and minimal guide on the process can be viewed by hovering the mouse over the process.

### Proofreading cell segmentation using nuclear segmentation

PlantSeg-tools offers this script for proofreading cell segmentation based on nuclei knowledge (https://github.com/hci-unihd/plant-seg-tools). The method is first described in this manuscript and is part of this study. The cell segmentation will be adjusted to resolve any conflict with the respective nuclear segmentation, thus the accuracy of the nuclei is extremely important. Errors in nuclear segmentation are propagated to cell segmentation. The script is composed of two different subroutines. One for correcting the split errors in cell segmentation and one for fixing the merge mistakes. The split routine checks for each cell whether two or more nuclei (measured as a percentage of the total cell volume) overlap with the cell segmentation by more than a user-defined “threshold-split (t-split)”. If the overlap is above the threshold, the script will use the nuclear segmentation as seed and split the cell using the seeded watershed algorithm. The merge routine checks for each nucleus whether two or more cells (measured as a percentage of the total nucleus volume) overlap a single nucleus segmentation by more than a user-defined “threshold-merge (t-merge)”. If the overlap is above the threshold, the script will merge the cells. The default thresholds provided are 66% for “t-split” and 33% for “t-merge”.

### Optimized workflow from imaging to segmentation of nuclei dataset

Obtaining confocal Z slices is achievable with a recommended xyz voxel size ranging from 0.12 x 0.12 x 0.25µm³ to 0.25 x 0.25 x 0.25 µm³, ensuring visually identifiable non-oversaturated nuclei signals. For optimal results, we propose imaging with line average ranging from 2 to 5 whenever feasible. Employing microscope objectives with a high numerical aperture (ideally around 1.2 NA or higher) is advised. Nevertheless, both the PlantSeg and the StarDist-ResNet platinum models are quite flexible to the imaging conditions as they were able to process a range of image quality (Table. 3). For nuclei segmentation using the two platinum models, we present GoNuclear (https://github.com/kreshuklab/go-nuclear). GoNuclear comes with the PlantSeg and StarDist-ResNet platinum models. Although the results are comparable, we recommend trying StarDist with the StarDist-ResNet platinum model first, as it is a bit less involved compared to the PlantSeg 3D nuclei segmentation pipeline. GoNuclear can batch process nuclei images and output segmentation can be saved as a tiff/HDF5 file which can be imported into MorphoGraphX. As an alternative, the PlantSeg_3Dnuc_platinum model has been integrated into MorphoGraphX, allowing 3D nuclear predictions to be generated, which can then be 3D segmented using the ITK watershed algorithm, all within MorphoGraphX. MorphoGraphX enables multiple 3D stacks and segmented images to be superimposed on each other, allowing the data sets to be proofread as needed. A 3D nuclei mesh can be created in MorphoGraphX and quantifications can be performed. Numerical results can be exported as a csv file for further processing.

## Acknowledgements

We acknowledge support by EMBL IT Services and the Center for Advanced Light Microscopy (CALM) of the TUM School of Life Sciences.

## Data availability

Information and code for training and inference using PlantSeg, Cellpose, or StarDist, including how to segment new 3D nuclei volumes as mentioned in this study, can be found in the GoNuclear repository: https://github.com/kreshuklab/go-nuclear. Other software can be downloaded at the following links: MorphoGraphX: https://morphographx.org. PlantSeg: https://github.com/hci-unihd/plant-seg. Plant-seg-tools: https://github.com/hci-unihd/plant-seg-tools. StarDist: https://github.com/stardist. Cellpose: https://github.com/mouseland/cellpose. We provide the 2 platinum models through the BioImage Model Zoo (https://bioimage.io) for FAIR use through different client tools of our community. PlantSeg_3Dnuc_platinum: Zenodo ID 0.5281/zenodo.8401064; Zoo name: efficient-chipmunk. StarDist3DResnet_3Dnuc_platinum: Zenodo ID: 10.5281/zenodo.8421755; Zoo name: modest-octopus. All datasets used for the figures and the entire bundle of models we trained can be downloaded from BioImage Archive (BIA) (https://www.ebi.ac.uk/bioimage-archive/) (Hartley et al., 2022)/BioStudies (https://www.ebi.ac.uk/biostudies/) (Sarkans et al., 2018), accession S-BIAD1026. The MorphoGraphX Process “Mesh/Nucleus” is available with version 2.0.2.and above https://morphographx.org. The data used for quantification of the Arabidopsis ovule N/C ratios include the training datasets generated in this study (Biostudies accession S-BIAD1026) and were also obtained from (Vijayan et al., 2021) (BioStudies, accession S-BSST475). The mouse embryo BlastoSPIM data set (Nunley et al., 2023) can be downloaded from the respective website (https://blastospim.flatironinstitute.org/html/series.html).

## Supplementary Materials

Attached as a separate file.

1. Supplementary Results

2. Supplementary File

